# The Biology of ‘Risk-On’. Decreasing Inflammatory and Stress Responses on a London Trading Floor

**DOI:** 10.1101/473157

**Authors:** N. Z. Xie, L. Page, D. A. Granger, J. M. Coates

## Abstract

Human physiological arousal is highly sensitive to information and uncertainty. Little is known, however, about how to measure information in natural settings, nor about which physiological systems respond to it. Financial market prices, and their volatility, present a convenient measure of informational load. Here we report on a study into the physiological response of traders in the City of London during a period of extreme, but declining, volatility. We sampled salivary cortisol, the main stress hormone, and the pro-inflammatory cytokines, IL-1*β*, IL-6, IL-8, and TNF-*α* three times a day for two weeks. We found that average daily cortisol levels tracked closely an index of equity and bond volatility, as did levels of IL-1*β*. Within-day cortisol and IL-1*β* levels also tracked one hour lagged volatility. Interestingly, the cascade of endocrine and immunological changes was initiated by IL-1*β*, the first responder of the stress and inflammatory responses. Our results have implications for finance because chronic stress and the immune response known as ‘sickness behaviour’ could have powerful effects on risk-taking and market stability.

## Introduction

Information exerts broad and powerful effects on human physiological arousal. Humans do not process information dispassionately, as computers do; we react to it physically. Upon receipt of a startling piece of news, for example, we experience an increased heart rate, faster breathing, dilated pupils, and a surge of adrenalin, cortisol, and pro-inflammatory cytokines. The information, in short, has stimulated a physical arousal known as the stress response.

The stress response is poorly named as it implies an aversive experience. In fact, the stress response is nothing more than a metabolic and cardiovascular preparation for potential physical movement. It involves recruiting the substrates for oxidative metabolism, i.e., glucose from glycogen stores in muscle and liver, and free fatty acids from lipids; as well as the activation of the cardiovascular system to distribute this fuel to stress relevant tissues throughout the body. Central to this metabolic preparation is the hormone cortisol, a glucocorticoid produced by the adrenal cortex. As levels of cortisol rise, especially if they segue from a short-lived acute elevation to a long-lived chronic one, they feedback on the brain altering attentional control^1^, memory recall ^2^ and risk-taking ^3 4^. The close linkage between information, physiology and cognition raises difficult questions about the possibility of studying cognitive activity without reference to accompanying somatic changes ^5^.

Cortisol levels have been found to increase in situations of novelty and uncertainty ^6 7 8^. These situations can be described as information rich. According to Claude Shannon, inventor of information theory, the amount of unpredictability in a series of events is a measure of its informational content, or entropy ^9^. A signal that is predictable carries no information, just as news that is already known is no longer news; while the result of a coin toss, because of its unpredictability, carries maximum information. We attend to information and ignore that which is predictable^10 11^, although as entropy increases we eventually approach pure noise, to which we quickly habituate. Some information theorists argue rather that we attend to information containing Bayesian surprise, i.e., a discrepancy between our prior and posterior distributions ^12^. When immersed in an information-rich environment, one where we are exposed to continuous novelty and surprise, we do not know what to expect and increasing cortisol levels help us marshal a preparatory stress response ^13^.

The close linkage between information and arousal can be observed vividly and measured objectively in the financial markets. The more information flowing into the markets the greater the volatility and the greater the uncertainty. For this reason one closely-followed index of volatility in US stocks, the VIX, has been aptly termed, ‘The Fear Index’. When traders and asset managers around the world read news feeds and observe price changes they become immersed in an environment of nonstop information and surprise, and their physiology tracks it in sympathy. In a previous study conducted in the City of London we observed that traders’ physiological arousal calibrated to informational load, with cortisol levels following market volatility higher over a two week period (R^2^=0.86) ^14^ In a follow-up laboratory study we replicated the observed increase in cortisol by administering hydrocortisone to volunteers, and found that the chronic elevation of cortisol decreased their risk appetite 44% ^4^ Here the physiological arousal was altering the participants’ decision-making and risk-taking.

These studies raised a question: what other physiological systems track the informational challenge of a volatile market? One system in particular seemed to us relevant for risk attitudes – the immune system. Repeated acute stress or prolonged stress can promote systemic and chronic inflammation and a phenomenon known as ‘sickness behaviour’, a condition of fever, anhedonia, social withdrawal, enhanced fear learning, and reduced exploratory behaviour, i.e., reduced risk-taking ^15 16 17^.

The first step in the cascade of events leading to sickness behaviour is the triggering of inflammation by the release of cytokines, the signalling proteins of the immune system. In the presence of a pathogen or infection, cytokines are released by cells of the innate immune system, such as macrophages and monocytes. Some cytokines, for example a subset known as chemokines, create a chemical gradient along which T cells and natural killer cells can navigate towards the site of infection or injury. Recent research has also found that pro-inflammatory cytokines are released during episodes of psychological stress, even in the absence of a pathoge ^18 19 20^, a process sometimes called ‘sterile inflammation’^21^. Upon investigation immunologists found that cytokines play an important metabolic role in the stress response ^22 23^.

Key to the inflammatory and metabolic cascade coordinated by cytokines is interleukin-1β (IL-1β), which acts as the ‘first responder’ of the immune system ^23 24^, stimulating production of the other pro-inflammatory cytokines (such as IL-6, IL-8, and TNF-*α*) ^25^. It is also the key cytokine promoting sickness behaviour ^15 26 27^, with other cytokines such as IL-6 amplifying its effects ^28 29^. During a stressful psychological challenge, IL-1β releases free fatty acids from adipocytes, as well as glycerol - providing the substrate for gluconeogenesis - and it promotes the uptake of glucose into stress-sensitive tissues in the brain and heart ^22^. Interestingly, IL-1β also acts as an upstream signalling molecule for the release of cortisol itself ^30 31^. It has been suggested that the endocrine system was built upon the evolutionarily older immune system and today makes use of the latter’s extensive and rapid surveillance network ^19^.

To further investigate the endocrine and immunological response of traders to informational load we sampled salivary cortisol and the pro-inflammatory cytokines IL-1*β*, IL-6, IL-8 and tumour necrosis factor (TNF-*α*) from a group of traders working at a hedge fund in the City of London. The study aimed in the first instance to ascertain if the remarkably high correlation between volatility and cortisol found in our previous study replicates. That first study was conducted during a period of rising volatility and increasing cortisol. The present study, on the contrary, was conducted towards the end of the European Sovereign Debt Crisis, a period of extremely high, yet declining volatility. This was also a time of increasing risk-taking across global markets– a process known colloquially in the financial world as ‘risk-on’. The timing allowed us to discover if declining volatility led to a declining stress response among the financial community.

Finally, the sampling of IL-1β, as well as other downstream pro-inflammatory cytokines, permitted us to discover if the immune system tracked information to the same extent as the endocrine. Together these physiological markers might indicate the biological changes occurring in the financial community during periods of risk-on.

## Methods

We recruited 15 male traders from a mid-sized hedge fund in the City of London. The traders varied in age from 21 to 39 years with a mean of 30.3 years. They bought and sold futures contracts, mostly in the equity and bond markets; or they executed spread trades, buying one contract and selling a different one, to profit from relative price movements. Their trade size could amount to **€** billions, but they held their positions for short periods of time, minutes or hours, only occasionally for days.

This style of trading benefits from short-term inefficiencies in asset prices and in delays between the release of news stories and their effect on markets. It therefore requires intensive scrutiny of news and price patterns, as well as the cognitive and physical speed to interpret news and execute a trade before other traders or computerized trading programs do so. To support the traders, the trading stations come equipped with real-time price feeds from futures exchanges; newsfeeds from major news outlets around the globe; a risk management system which displays the traders’ ongoing profits and losses (known as P&L) and their risk (known as value at risk, or VaR); and an intercom system over which a resident economist comments on the news and the economic statistics being released in the major economies. These traders, in short, work in an environment that is information-rich, and they deal constantly with surprise.

We conducted our study during a particularly volatile period in European markets, the tail end of the European Sovereign Debt Crisis. We monitored these traders onsite for two weeks – 10 business days - taking saliva samples via passive drool three times a day, at 9am, 12pm, and 4pm UK time. Salivary cortisol correlates reliably with serum, largely because steroids from the blood infiltrate saliva by passive diffusion; but research using salivary cytokines is relatively new. Some studies have successfully employed salivary cytokines, and have found a significant correlation between them and serum measures ^25 32 33 34^, especially in the case of IL-1ß ^35 36^. Cytokines, being proteins, do not infiltrate easily from blood to saliva; but it has been suggested that during stress salivary glands are stimulated by the sympathetic nervous system to produce local inflammation, this reaction comprising one part of a more systemic mucosal immune response. In this manner, a local salivary response may provide a window into systemic inflammation ^25^.

Samples were packed in ice for initial transport, temporarily stored at −80C, and then packed in dry ice for transport to the Institute for Interdisciplinary Salivary Bioscience Research. All samples arrived frozen. They were assayed for cortisol, IL-1ß, IL-6, IL-8, and TNF-*α*. Cortisol was analysed using a commercially available competitive immunoassay without modification to the manufacturer’s (Salimetrics, Carlsbad, CA) recommended protocol. Cytokines were measured using a 96-well multiplex (4-plex) electrochemiluminescence immunoassay from MesoScale Discovery (MSD, Gaithersburg, MD). Each well of a plate was coated with capture-antibodies specific to IL-1β, IL-6, IL-8, and TNFα. Detection antibodies were coupled to SULFOTAGTM labels that emit light when electrochemically stimulated via carbon-coated electrodes in the bottom each microwell. The assay was run using standard diluent (MSD # R51BB). Cytokine concentrations (pg/mL) were determined with MSD Discovery Workbench Software (v. 3.0.17) using curve fit models (4-PL with a weighting function option of 1/y2). All lower limits of detection were below 0.50 pg/mL, average intra-assay coefficients of variation (CVs) were less than 5%, and average inter-assay CVs were less than 10%.

We recorded all the markets traded by the traders during the study. The traders traded a range of futures contracts in the bond, equity, currency, and commodity markets, but their largest risk exposure was to bonds and to equities (see Supp Info, Table 1). Within these sectors their specific exposure was mostly to German bond futures, known as Bunds, and to a European equity index, known as the EuroStoxx 50. We therefore constructed an index of the markets they traded, weighted by number of trades, consisting of 66.9% Bunds and 33.1% Eurostoxx. Adding further securities did not affect our statistical analysis, either because the further securities were highly correlated with Bunds or Stoxx, or because adding them had minimal effect on our results. The resulting index is a weighted average of the historical volatilities of these two securities:

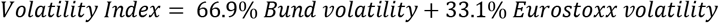

For both Bunds and Stoxx we downloaded futures exchanges’ official end-of-day prices, as well as historic volatilities for the previous 5, 10, and 30 day periods. Historic volatilities capture the actual variance of a market over a period of time, in our case 5, 10, and 30 days. We then combined volatilities for Bunds and Stoxx into our volatility index. Using it as a measure of the uncertainty faced by the traders over these different time periods, we were able to observe the effect of varying levels of uncertainty on the traders’ endocrine and immune systems. For data analysis, cortisol and cytokine measures were logarithm transformed and modelled by simple linear regression.

The study was conducted according to the code of ethics on human experimentation established by the Helsinki declaration and in accordance with the approved guidelines. The study was approved by the ethics committee of the School of Biological Sciences at the University of Cambridge. All participants provided written informed consent.

## Results

### Salivary cortisol and cytokines display the expected diurnal pattern

To begin our analysis we looked for evidence that our salivary cortisol and cytokine measures were behaving as predicted by existing literature. Cortisol levels are known to display a declining diurnal pattern, peaking prior to a person’s waking and then declining over the course of the day; and our cohort did display this pattern (Fig.1). Analysing data from 3 daily time points over ten days we found that a time dummy variable had a strongly negative coefficient when regressed against cortisol levels (*R*^2^=0.836, *p*<0.001, *N*=30).

**Figure 1.**
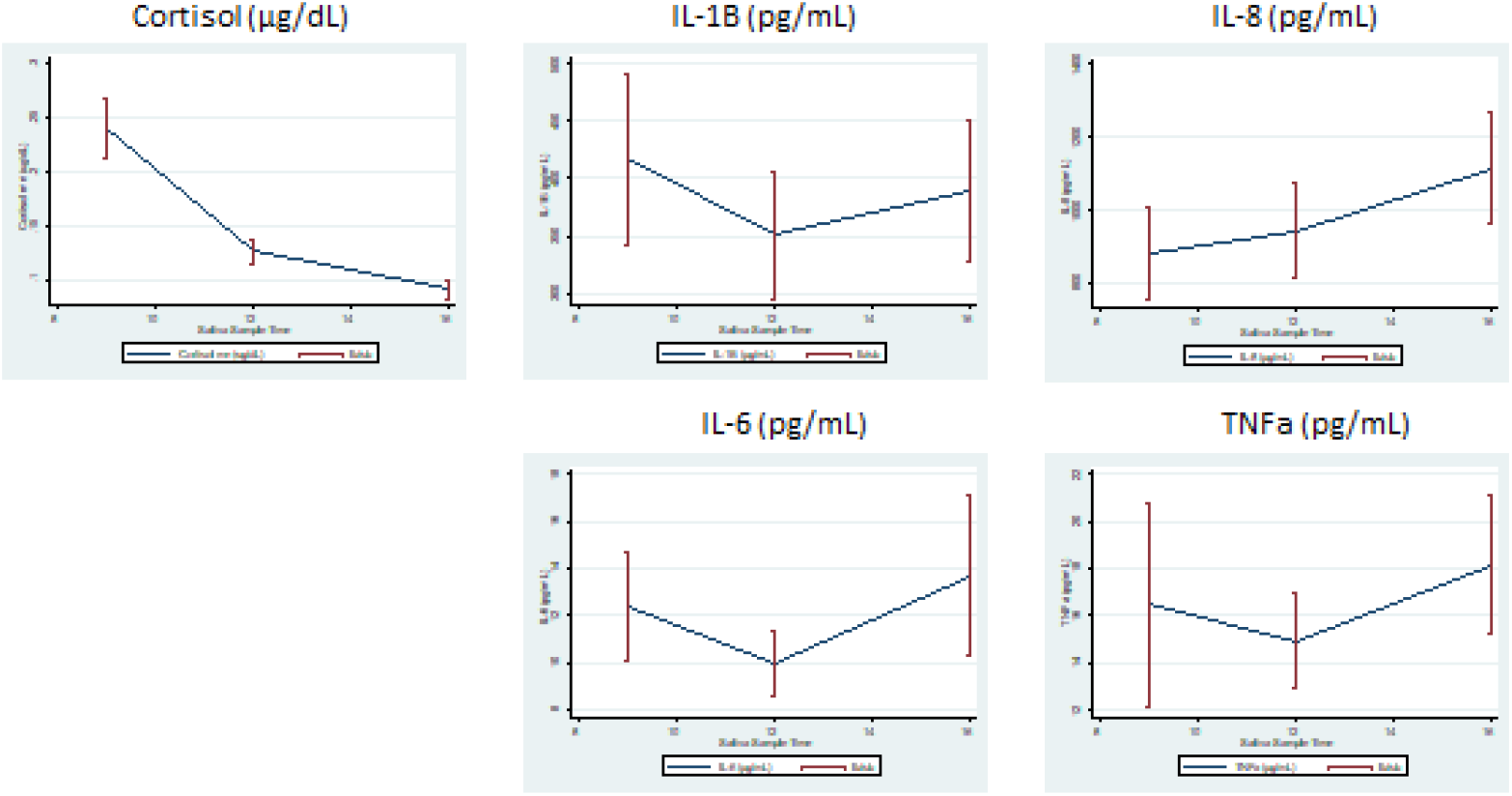
Diurnal pattern of cortisol and pro-inflammatory cytokines averaged from all traders at 9am, 12pm, 4pm. Traders’ cortisol levels over the course of the day display the normal downward slope. Cytokines display rising levels – albeit non-significantly - over the course of the day, as the anti-inflammatory effects of cortisol decline.

Cytokine levels commonly display the opposite diurnal pattern to cortisol, either rising over the course of the day ^37^, or dropping between early morning and noon, and rising thereafter to peak late at night ^38^. The reason advanced for this cytokine pattern is that as cortisol levels fall during the day its powerful anti-inflammatory effects decline, leading to higher pro-inflammatory cytokine levels. We did observe this pattern in our cohort (Fig.1), although the time dummy when regressed against cytokine levels was significant only for IL-8 (data not shown). Existing literature has also found that changes in IL-1ß lead the other pro-inflammatory cytokines ^23 25^, and we too replicated this result, as reported below. These results suggest that our salivary measures were well behaved.

### Daily average cortisol levels track volatility

To test the hypothesis that traders’ stress responses track market uncertainty, we compared daily average cortisol levels (averaged from 3 time points) for all 15 traders with historic volatility. During the two weeks of the study our volatility index dropped approximately 18%, and average daily cortisol levels dropped 19% (Fig.2a). We regressed these cortisol levels against 5, 10, and 30 day historic volatilities of our Bund and Eurostoxx index and found that the statistical results varied systematically and consistently across time horizons (Fig.2a). The strongest effects of volatility on cortisol occurred at the 10 day horizon, with a positive and highly significant correlation between volatility and cortisol (coef. = 0.093, Adj *R*^2^ = 0.688, *p* =0.003, *N* = 10). The effect of volatility on cortisol appears to increase from the 5-day horizon to the 10-day and to decline thereafter: there was a low and non-significant correlation between cortisol and 5-day volatility, rising to the 10-day horizon, then decaying, with 30-day volatility showing a positive and significant correlation, yet weaker than the 10-day (Table 2).

**Figure 2.**
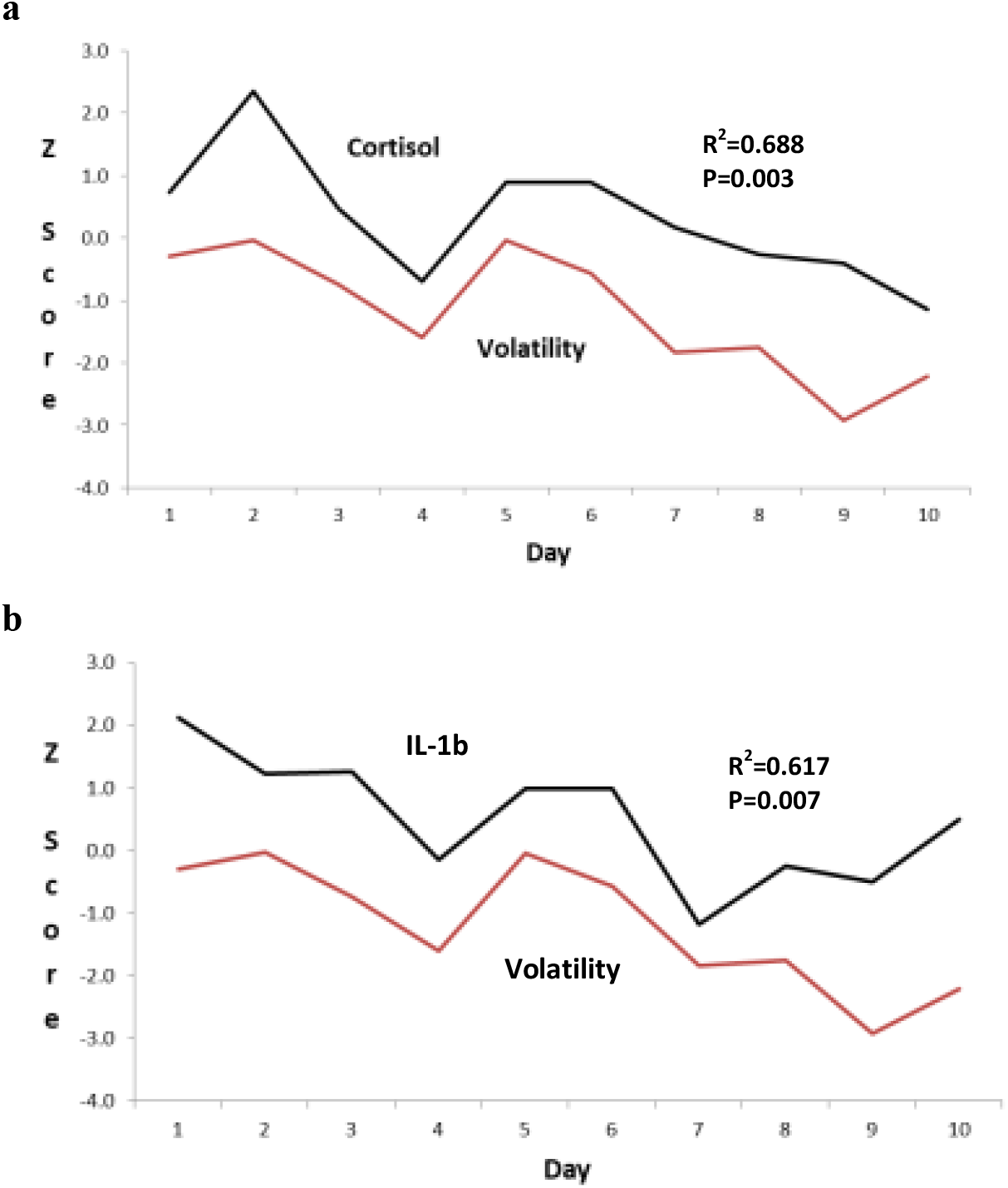
Cortisol and IL-1b plotted against 10-day historic volatility. **a)** Upper curve is plot of cortisol levels averaged from all traders for each day of the 10 day study. Lower curve is volatility or variance of our market index (consisting of European bonds and stocks) calculated over the previous 10 days. **b)** Same curves for 1L-1*β*. Z-scores in figures have been adjusted by adding or subtracting a constant in order to prevent the two curves from lying on top of each other.

**Table 2.**
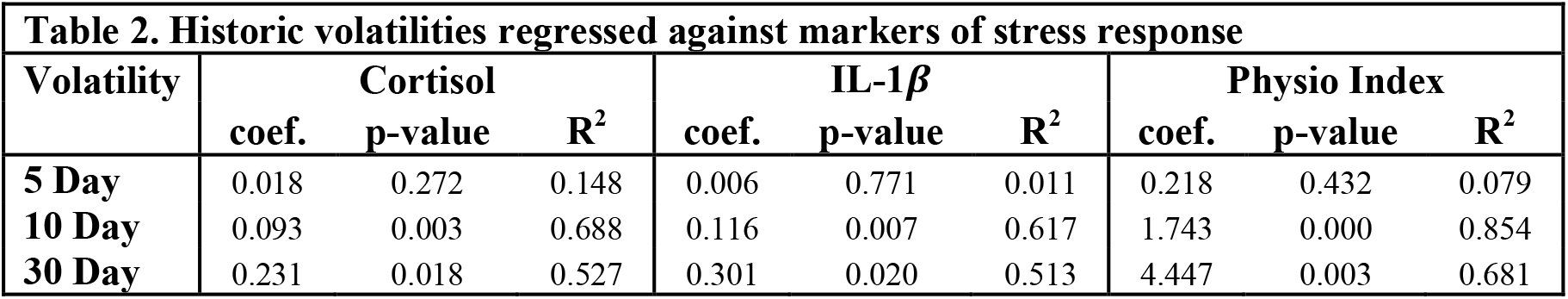
Correlations by time horizon. Correlations between volatility and cortisol, IL-1b, and the physiological arousal index at the 5, 10, 30 day time horizons. All three markers of physiological arousal display a consistent pattern: correlations between arousal and 5 day volatility are positive but weak, rise to a maximum at the 10 day horizon; and decay out to the 30 day horizon.

| Table 2. Historic volatilities regressed against markers of stress response |  |  |  |  |  |  |  |  |  |
| --- | --- | --- | --- | --- | --- | --- | --- | --- | --- |
| Volatility | Cortisol | | | IL-1β | | | Physio Index | | |
|  | coef. | p-value | R<sup>2</sup> | coef. | p-value | R<sup>2</sup> | coef. | p-value | R<sup>2</sup> |
| 5 Day | 0.018 | 0.272 | 0.148 | 0.006 | 0.771 | 0.011 | 0.218 | 0.432 | 0.079 |
| 10 Day | 0.093 | 0.003 | 0.688 | 0.116 | 0.007 | 0.617 | 1.743 | 0.000 | 0.854 |
| 30 Day | 0.231 | 0.018 | 0.527 | 0.301 | 0.020 | 0.513 | 4.447 | 0.003 | 0.681 |

### Daily average IL-1β levels track volatility

Similar results were found when we looked at the immune response, as proxied by IL-1ß levels, to changes in uncertainty. Over the course of the study average daily IL-1ß levels fell 18% (Fig.2b). Regressing IL-1ß against our volatility index again showed a positive but non-significant correlation over 5 days; a positive and highly significant correlation over 10 days (coef. = 0.116, Adj *R*^2^ = 0.617, *p* = 0.007, *N* = 10); and a significant but weaker correlation over 30 days (Table 2).

### Index of physiological arousal tracks volatility

As cortisol and IL-1*β* are both molecules involved in the stress response we next looked to see how they jointly responded to uncertainty. We combined Z-scores for cortisol and IL-1*β* into an index of physiological arousal. When we regressed this index of arousal against volatility we found even higher correlations than for either marker on its own, and the same pattern across time horizons, with a weak correlation over 5 days; a very high correlation over 10 days (coef. = 1.743, Adj *R*^2^ = 0.854, *p* < 0.001, *N* = 10); and a decaying influence over 30 days (Table 2).

### Within-day cortisol and IL-1*β* levels track 1 hour lagged volatility

The results for our daily averages demonstrated gradual fluctuations in baseline cortisol and IL-1*β* levels over the two weeks of the study. We were also able to observe more rapid, within-day dynamics because we had access to minute-by-minute prices for both Bunds and Eurostoxx from which we could calculate within-day volatilities. We calculated 1 hour lagged volatility for our index of Bunds and Eurostoxx, and then regressed this against cortisol and IL-1*β* levels at each of our three sampling times. As both cortisol and IL-1*β* display strong diurnal patterns we controlled for these changes by adding to our regression equation a dummy variable for sampling time.

We found that the average cortisol level at each sampling point was significantly and positively correlated with 1-hour lagged variance of our index of Bunds and EuroStoxx (coef. = 0.182, Adj *R*^2^ = 0.86, *p* =0.02, *N* = 30). This finding agrees with existing research suggesting that the half-life of an episodic increase in cortisol is approximately 66 minutes ^39^. We found similar results when regressing 1-hour variance against IL-1*β* (coef. = 0.276, Adj *R*^2^ = 0.23, *p* =0.01, *N* = 30), and our index of physiological arousal (coef. = 1.51, Adj *R*^2^ = 0.60, *p* =0.002, *N* = 30).

### IL-1β levels predict those of cortisol and pro-inflammatory cytokines

Finally, we investigated the interactions between cortisol and the four pro-inflammatory cytokines. We employed the vector autoregressive (VAR) model to capture the linear inter-temporal dependencies among cortisol, IL-1*β*, IL-6, IL-8 and TNF-α. We then applied the Granger causality Wald test to test whether prior values of cortisol and cytokines could predict future values of the other molecules. We used the previous day (*t* – 1) averaged responses to predict the values on day *t*. For VAR equations, estimation results, and Granger causality test results are presented in Table 2 and 3 in Supplementary Information. The time series of the cytokines were checked for stationarity by the Dicky-Fuller unit root test, to meet the prerequisite of applying the Vector Autoregressive model.

We found that IL-1*β* significantly predicted the other cytokine levels as well as cortisol indirectly (Fig. 3) a finding that is broadly consistent with previous research finding that IL-1*β* is an upstream signalling molecule for other inflammatory processes and for the stress response^25 30 31^. We should however point out the small sample size underlying this analysis.

**Figure 3:**
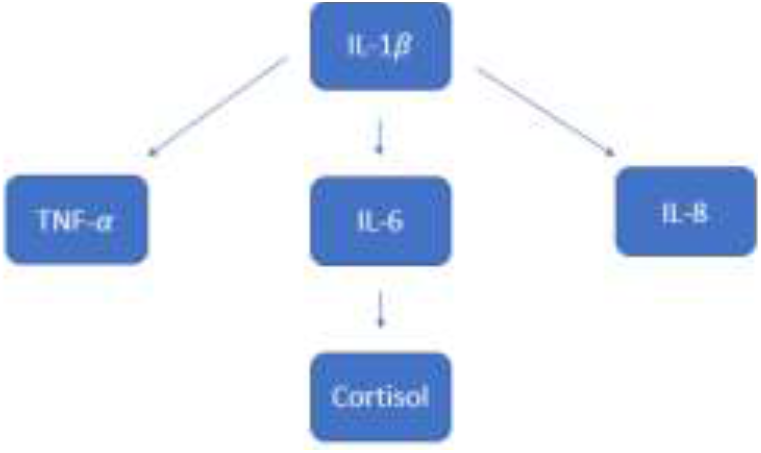
Granger causality Wald test results. The arrows represent the direction of significant causalities.

## Discussion

The present study was conceived as field work ^40^, specifically to observe the physiological arousal occurring in traders as informational load varied. Fluctuating securities prices, the release of economic statistics, the reporting of news stories – all these inject information into the markets. The greater the amount of information and the greater the amount of surprise contained in it, ^10 12^ the greater the ongoing uncertainty, and the higher the market volatility. Uncertainty triggers in traders and in the financial community more generally a preparatory physical arousal in the form of the stress response even though the stressor is psychological and most employees of financial firms sit at desks.

We were able to observe this phenomenon in real traders while they traded the markets. We found that daily average cortisol levels tracked closely an index of volatility calculated for the previous 10 days. The same result was found for IL-1*β* (Fig. 2). As IL-1*β* acts as an upstream signal in the stress response and works in concert with cortisol, we combined cortisol and IL-1*β* into an index of physiological arousal and found it tracked volatility even closer. In this study we had three daily sampling times, permitting us to look at intra-day dynamics, and here we found that cortisol, IL-1*β*and our index of arousal all tracked variance of the securities index over the previous hour. Finally, using Granger causality analysis, we found, as has previous research, that IL-1β predicted subsequent changes in cortisol and the other pro-inflammatory cytokines, IL-6, IL-8, TNF-*α* (Fig. 3).

The various time horizons over which we measured volatility provide some insight into the length of time over which uncertainty must persist in order to alter baseline cortisol and cytokine levels. 5-day volatility showed a positive yet weak and non-significant correlation with cortisol, IL-1*β*, and our index of physiological arousal. The correlations rise strongly to the 10-day horizon, and decay after that (Table 2). Related to these findings are those from a study of cortisol and risk preferences that administered hydrocortisone to volunteers over an 8-day period and found that acute, 1-day changes in cortisol levels had no significant effect on risk preferences but that an 8-day exposure did. The data suggests that the 8 to 10-day horizon may be key in causing chronic changes to arousal.

These results could have implications for economics. Many branches of economics must estimate the time horizon over which a person’s expectations are formed, a process known as ‘adaptive’ or ‘lagged’ expectations^41^. These expectations may relate to future inflation, financial returns, or volatility of the markets. Physiology may assist in the debates surrounding expectations by identifying the time periods over which baseline physiological arousal alters, and with it our risk preferences.

This study was also intended as part of an ongoing research program into the causes of financial market instability. Our guiding hypothesis has been that physiological changes occurring in the financial community shift risk preferences pro-cyclically, causing investors to take too much risk on the upside, pushing a bull market into a bubble, and too little on the downside, driving a bear market into a crash ^5 13^.

Key to this risk-taking behaviour is the difference between an acute and chronic stress response. An acute increase in cortisol marshals metabolic reserves and has been found to aid in the recall of important memories^42^, to promote motivated behaviour^43^, but to leave financial risk preferences largely unchanged ^4^. Chronic stress, on the other hand, can have opposite effects. By reducing spine density in the hippocampus^44^ and causing dendritic arborisation in the amygdala^45^, chronically elevated cortisol levels can promote anxiety and depression ^46^, impair behavioural flexibility^47^, and instil risk-aversion^48^. An earlier study found that cortisol on a trading floor rose 68% over a two week period as volatility increased ^14^; and a subsequent lab-based study found that a cortisol response of this size and duration caused a 44% increase in risk-aversion ^4^. Shifts in risk preferences of this magnitude could well destabilize markets.

A similar pattern exists for acute and chronic inflammatory responses ^49^. An acute stressor enhances innate and adaptive immunity and through the actions of the sympathetic nervous system triggers the production of pro-inflammatory cytokines, most notably IL-1β ^50 51^. This cytokine response can occur during both exercise and psychological stress, even in the absence of a pathogen. The stress response is largely a metabolic preparation for movement and IL-1β enhances glucose uptake, in an insulin independent manner, into stress relevant tissues such as brain, heart, and skeletal muscle ^22 23^.

It also exerts central effects^52^. IL-1*β* does not cross the blood brain barrier, but peripherally produced IL-1β can nonetheless gain access to the brain via the circumventricular organs where the blood brain barrier is weak; via active transport of smaller molecules such as prostaglandins^53^; or via innervation of the vagus nerve which leads to the central production of IL-1β ^54 55^. Receptors for IL-1β are found in the paraventricular nucleus of the hypothalamus and in the pituitary and it is from here that IL-1β stimulates the production of CRH, ACTH and cortisol^56^. Receptors are found as well in the central amygdala and the hippocampus; and in the latter, during acute stress, IL-1β aids in the formation of memories^57 58^.

Chronic stress, on the other hand, can trigger low grade systemic inflammation and impair memory formation ^59 60^. It also produces sickness behaviour, a condition of fever, anhedonia, enhanced fear learning, social withdrawal and, if prolonged, depression ^15 17^. Sickness behaviour also features reduced locomotor activity and exploratory behaviour ^61^, which implies an element of risk-aversion. An extended period of low grade inflammation and sickness behaviour could therefore augment the effects of chronic cortisol exposure by instilling risk-aversion among the financial community.

Indeed, chronic inflammation induced risk-aversion could well provide an explanation for the observation that stock markets under-perform during the time of year when seasonal affective disorder occurs ^62 63^. It is at this time of year, late autumn, early winter (October is a notorious month for market crashes) that circulating cytokines, including IL-1β and TNF-α and IL-6, tend to increase ^64 65^.

We conducted our study towards the end of the European Sovereign Debt crisis, when volatility was decreasing from elevated levels. During this chapter of the crisis, dominated by the Greek bailout negotiations, volatility on Stoxx rose from 19% in July 2011 to 48% in Sept, and then decayed to 24% by the end of our study. During the study average daily cortisol levels fell 19% and IL-1β 18% (Fig.2). We cannot know the levels these molecules reached during the crisis, nor for how long; but given the high correlations we have found in this and our previous study, we suggest that both cortisol and IL-1β rose during the Debt Crisis, perhaps substantially, and for an extended period of time. This chronic elevation would have brought risk-aversion to the financial community and may well have exacerbated the market sell-off.

When we came onto the trading floor the crisis was abating, and the traders’ physiology may have been returning to baseline. Decreasing stress and inflammatory responses could have released the brake on their risk-taking and they, along with the wider financial community, felt emboldened to start buying in the credit markets. Our findings therefore offer the beginnings of an understanding of what could be called the biochemistry of risk-on, and more generally of the shifting risk preferences that destabilize markets ^5^.

Finally, our findings also carry practical implications as they suggest that uncertainty could be used as a little-appreciated tool of monetary policy. Central banks today are increasingly taking responsibility for dampening bull markets before they become bubbles and pose a risk to financial stability. The conundrum central bankers face is that if they reign in market exuberance by raising interest rates they may inadvertently harm the economy. However, they could solve this conundrum if they appreciated that they have in their policy toolbox not just the level of interest rates but also the variance of rates as well as uncertainty surrounding their intentions. Increasing uncertainty surrounding future interest rates could inject risk-aversion into the markets and calm a bull market before it morphs into a bubble.

Paul Volker, a former chairman of the US Federal Reserve, was a master of using uncertainty as a policy tool; but subsequent chairmen, beginning in the early 1990s, have adopted a policy known as ‘forward guidance’ which involves communicating clearly to the markets the central bank’s intentions. This well-meaning policy, by reducing uncertainty, may have inadvertently contributed since the advent of forward guidance to the increased violence of market bubbles and crashes, such as the Asian Financial Crisis of 1998, the Dot.com Bubble, and the Financial Crisis of 2008-09. Increasing the variance of short rates, by instilling risk-aversion among the financial community through the mechanisms outlined above, could help reign in irrational exuberance before it becomes dangerous.

## Supporting information

## Acknowledgements

We thank Narayanan Kandasamy and Mark Gurnell of Dept of Medicine, University of Cambridge for assistance with the study; and Bruce McEwen, Robert Sapolsky, and Christopher Coe for comments on manuscript. Any mistakes are ours, not theirs. J.M.C. was supported by a Programme Grant from the UK Economic and Social Research Council.

## Author Contributions

J.M.C. conducted study; N.Z.X., L.P., D.G. and J.M.C. analysed data; J.M.C. wrote paper; all co-authors commented on manuscript.

