## Supplementary material for "The Biology of ‘Risk-On’. Decreasing Inflammatory and Stress Responses on a London Trading Floor"

**Table 1. Traded Markets: % of Total Trades**

| Bonds & Interest Rates | | | | | | Equities | | | Commodity | | FX |
|---|---|---|---|---|---|---|---|---|---|---|---|
| Europe | | | | UK | US | Europe | | US | | | |
| Bund | Bobl | Schatz | Euribor | Gilt | Notes | Stoxx | Dax | S&P | Gold | Oil | $/ € |
| 32.2% | 4.7% | 3.0% | 3.6% | 11.8% | 8.0% | 16% | 6.2% | 5.0% | 2.7% | 4.1% | 2.7% |
| 43.5% | | | | 11.8% | 8.0% | 22.2% | | 5.0% | 6.8% | | 2.7% |
| 63.1% | | | | | | 27.4% | | | 6.8% | | 2.7% |

**Future Contracts Legend**. Bund: German 10yr bond. Bobl: German 5yr bond. Schatz: German 2yr bond. Euribor: Euro 3 month deposit rate. Gilt: UK 10yr bond. Notes: US 10yr bond. Stoxx: Eurostoxx equity index. Dax: German equity index. S&P: U.S. equity index.

The index of the markets traded was weighted by number of trades, consisting of 66.9% Bunds (32.2%/(32.2%+16%)) and 33.1% Eurostoxx (16%/(32.2%+16%))

**Vector Autoregressive (VAR) models**.

$$\begin{cases} Cortisol_t = \beta_{10} + \beta_{11}Cortisol_{t-1} + \beta_{12}IL1\beta_{t-1} + \beta_{13}IL6_{t-1} + \beta_{14}IL8_{t-1} + \beta_{15}TNF\alpha_{t-1} + \varepsilon_{1t} \\ IL1\beta_t = \beta_{20} + \beta_{21}Cortisol_{t-1} + \beta_{22}IL1\beta_{t-1} + \beta_{23}IL6_{t-1} + \beta_{24}IL8_{t-1} + \beta_{25}TNF\alpha_{t-1} + \varepsilon_{2t} \\ IL6_t = \beta_{30} + \beta_{31}Cortisol_{t-1} + \beta_{32}IL1\beta_{t-1} + \beta_{33}IL6_{t-1} + \beta_{34}IL8_{t-1} + \beta_{35}TNF\alpha_{t-1} + \varepsilon_{3t} \\ IL8_t = \beta_{40} + \beta_{41}Cortisol_{t-1} + \beta_{42}IL1\beta_{t-1} + \beta_{43}IL6_{t-1} + \beta_{44}IL8_{t-1} + \beta_{45}TNF\alpha_{t-1} + \varepsilon_{4t} \\ TNF\alpha_t = \beta_{50} + \beta_{51}Cortisol_{t-1} + \beta_{52}IL1\beta_{t-1} + \beta_{53}IL6_{t-1} + \beta_{54}IL8_{t-1} + \beta_{55}TNF\alpha_{t-1} + \varepsilon_{5t} \end{cases}$$

Table 2: VAR Model Estimation Results:

| | (1) | (2) | (3) | (4) | (5) |
|---|---|---|---|---|---|
| | $Cortisol_t$ | $IL1\beta_t$ | $IL6_t$ | $IL8_t$ | $TNF\alpha_t$ |
| $Cortisol_{t-1}$ | -0.364*<br>(-1.78) | -0.108<br>(-0.73) | 0.065<br>(0.50) | 0.200<br>(1.45) | 0.123<br>(0.89) |
| $IL1\beta_{t-1}$ | 0.367<br>(1.02) | -0.162<br>(-0.63) | -0.752***<br>(-3.29) | -0.635***<br>(-2.63) | -0.783***<br>(-3.23) |
| $IL6_{t-1}$ | 0.825**<br>(2.01) | 0.218<br>(0.73) | 0.010<br>(0.04) | -0.084<br>(-0.31) | -0.081<br>(-0.29) |
| $IL8_{t-1}$ | 0.187<br>(0.35) | 0.164<br>(0.42) | 0.613*<br>(1.79) | 0.534<br>(1.47) | 0.659*<br>(1.81) |
| $TNF\alpha_{t-1}$ | -0.514<br>(-1.04) | -0.145<br>(-0.41) | 0.149<br>(0.47) | -0.091<br>(-0.27) | 0.003<br>(0.01) |
| Constant | 5.814*<br>(1.86) | 6.383***<br>(2.82) | 1.778<br>(0.89) | 5.955***<br>(2.83) | 2.181<br>(1.03) |

$N$ = 29. $z$ statistics in parentheses; *: $p < 0.1$, ** $p < 0.05$, *** $p < 0.01$.

Granger causality Wald test results:

Table 3: Granger causality Wald test results

| Equation | Excluded | $\chi^2$ | Degree of Freedom | $p$ value |
|---|---|---|---|---|
| Cortisol | IL-1$\beta$ | 1.048 | 1 | 0.306 |
| Cortisol ← | IL-6 | 4.058 | 1 | 0.044 |
| Cortisol | IL-8 | 0.120 | 1 | 0.729 |
| Cortisol | TNF-$\alpha$ | 1.084 | 1 | 0.298 |
| Cortisol | All | 11.603 | 4 | 0.021 |
| IL-1$\beta$ | Cortisol | 0.532 | 1 | 0.466 |
| IL-1$\beta$ | IL-6 | 0.539 | 1 | 0.463 |
| IL-1$\beta$ | IL-8 | 0.178 | 1 | 0.673 |
| IL-1$\beta$ | TNF-$\alpha$ | 0.164 | 1 | 0.685 |
| IL-1$\beta$ | All | 3.303 | 4 | 0.508 |
| IL-6 | Cortisol | 0.253 | 1 | 0.615 |
| IL-6 ← | IL-1$\beta$ | 10.850 | 1 | 0.001 |
| IL-6 | IL-8 | 3.195 | 1 | 0.074 |
| IL-6 | TNF-$\alpha$ | 0.225 | 1 | 0.635 |
| IL-6 | All | 16.054 | 4 | 0.003 |
| IL-8 | Cortisol | 2.116 | 1 | 0.146 |
| IL-8 ← | IL-1$\beta$ | 6.925 | 1 | 0.009 |
| IL-8 | IL-6 | 0.094 | 1 | 0.759 |
| IL-8 | TNF-$\alpha$ | 0.075 | 1 | 0.784 |
| IL-8 | All | 7.647 | 4 | 0.105 |
| TNF-$\alpha$ | Cortisol | 0.792 | 1 | 0.373 |
| TNF-$\alpha$ ← | IL-1$\beta$ | 10.429 | 1 | 0.001 |
| TNF-$\alpha$ | IL-6 | 0.085 | 1 | 0.771 |
| TNF-$\alpha$ | IL-8 | 3.280 | 1 | 0.070 |
| TNF-$\alpha$ | All | 12.820 | 4 | 0.012 |

Note: ← represents a significant test results ($p < 0.05$).
